# Benchmarking the Intratumoral Microbiome in Pancreatic Ductal Adenocarcinoma: A Longitudinal Assessment of Contamination Sources and Decontamination Strategies

**DOI:** 10.64898/2026.08.24.746744

**Authors:** Linh Dang, Louisa Eskelson, Jacob Hamm, Jutta Blumberg, Ulrike Wegener, Tim Beißbarth, Volker Ellenrieder, Albrecht Neesse, Christoph Ammer-Herrmenau

## Abstract

Pancreatic ductal adenocarcinoma (PDAC) harbors a distinct intratumoral microbiome. Yet rigorous characterization of its composition is hampered by pervasive environmental and procedural contamination. Sources of contamination have not been thoroughly explored, and the methods of decontamination have not been sufficiently evaluated in a benchmarking manner. We systematically collected >300 negative control (NCT) samples comprising paraffin from formalin-fixed paraffin-embedded (FFPE) samples, lysis buffer and sterile water over a period of four years processed by different laboratory persons (LP). All samples were sequenced using full-length 16S rRNA gene sequencing with Oxford-Nanopore Technologies. We benchmarked four decontamination methods (restrictive filtering, decontam, SCRuB, and the Nejman et al.-derived (Nj) pipeline) against fresh-frozen tumor samples (FF) from *LSL-Kras^G12D/+^*;*LSL-Trp53^R172H/+^*;*Pdx-1-Cre* (KPC) mice, using the abovementioned contamination assessment to calculate a composite score for the assessment. Further, we validated those methods via technical replicates. Microbial profiles of NCT samples were significantly determined by control type, LP, year and season reflecting complex batch effects. The 15 most abundant contaminants spanned well-characterized environmental taxa and human commensals from the oral cavity. The LP processing samples left a significant microbial trace highly contributing to the batch effect. Decontamination benchmarking demonstrated that the Nj method consistently outperformed alternatives in both composite score and inter-replicate concordance. Application of Nj to fresh frozen PDAC samples substantially reduced contaminant burden while preserving putative tumor-associated signals in FF but not FFPE samples. Our results support the adoption of the Nj decontamination approach for future intratumoral microbiome studies in fresh frozen tumor samples.

**Importance:** Tumors are increasingly recognized to harbor their own bacterial communities. For pancreatic ductal adenocarcinoma (PDAC)—one of the deadliest cancers—it has previously been shown that the intratumoral microbiome influences cancer proliferation and treatment response. But tumor tissue contains very little bacterial DNA, so trace contamination from reagents, lab surfaces, and the people processing samples can easily be mistaken for real biological signal. Using one of the largest longitudinal collections of negative control samples, together with tumor samples from a mouse model of pancreatic cancer, we show that the person handling a sample can introduce more bacterial noise than the sample’s environment, and we identify a decontamination strategy that reliably distinguishes genuine tumor-associated bacteria from background contamination in fresh-frozen tissue, but not in formalin-preserved tissue. This provides practical guidance for researchers designing low-biomass microbiome studies.

## Introduction

The intratumoral microbiome has emerged as a potentially important component of the tumor microenvironment, influencing immune evasion, drug metabolism, and clinical outcomes across multiple cancer types (1). To this end, the microbiome has been recently considered as one hallmark of cancer (2). In pancreatic ductal adenocarcinoma (PDAC), one of the most lethal malignancies, recent studies have reported the presence of distinct intratumoral bacterial communities that may contribute to disease pathogenesis and immunosuppression (3–5). However, a fundamental challenge in tumoral microbiome research is distinguishing genuine tumor resident microorganisms from environmental contaminants introduced during sample collection, processing, and sequencing. Low biomass specimens such as tumor samples are particularly vulnerable to contamination, as trace microbial DNA from reagents, laboratory surfaces, and laboratory personnel (LP) can outnumber endogenous taxa (6). Prominent environmental taxa such as *Sphingomonas*, *Ralstonia*, and *Pseudomonas* are routinely detected in negative controls and have been reported as dominant species in tumor microbiome studies that lack rigorous decontamination (3, 5, 7).

Several computational decontamination approaches have been proposed to address the contamination problem, including decontam (8), SCRuB (9), and the strategy employed by Nejman et al. (Nj) (6). These methods differ in their statistical assumptions, required inputs and aggressiveness of taxa removal. Nevertheless, a systematic, empirical comparison of their performance in the context of PDAC tumor microbiome profiling has been lacking. Here, we present a benchmarking framework that leverages a longitudinal collection of negative control (NCT) samples acquired across wet lab processing steps, combined with technical replicates of PDAC tissue samples. We evaluate four decontamination strategies using a novel composite score, balancing taxon yield and purity relative to the NCT-derived contaminant profile, and orthogonally validate findings using inter-replicate Aitchison distances. By applying full-length 16S rRNA gene sequencing with the Oxford Nanopore Technologies (ONT) platform we are able to perform microbial analysis on species level (10, 11). This work provides practical guidance for microbiome researchers seeking to generate reliable intratumoral microbial profiles from low-biomass PDAC specimens.

## Results

### 1. A longitudinal contamination assessment reveals a lab-specific contamination profile

#### 1.1 Different negative control types harbor distinct contaminant profiles

To comprehensively characterize the contaminant landscape, we collected 337 NCT at three stages of the wet-lab processing pipeline (Figure 1A), including 78 blank paraffin scratches (paraffin) from formalin-fixed paraffin-embedded samples (FFPE), 207 lysis buffer controls (buffer), and 52 sterile water samples used for diluting PCR-reactions (PCR), yielding an initial pool of 6,017 taxa. To reduce noise, we applied the following filtering steps: (i) samples with fewer than 500 reads were removed from further analyses; and (ii) extremely rare taxa were excluded using the following criteria: a detection threshold of at least 2 reads per sample, presence in at least one sample, and a total abundance of at least 1 × 10⁻⁶ across all samples, corresponding to 18 reads in our dataset. After filtering, 262 NCT remained: 75 paraffin, 167 buffer and 20 PCR controls, encompassing 3,073 taxa. These controls were accumulated between 2021 and 2025 (Table S1). Relevant batch covariates included were processing laboratory personnel (LP), season and year (Figure 1A).

**Figure 1:**
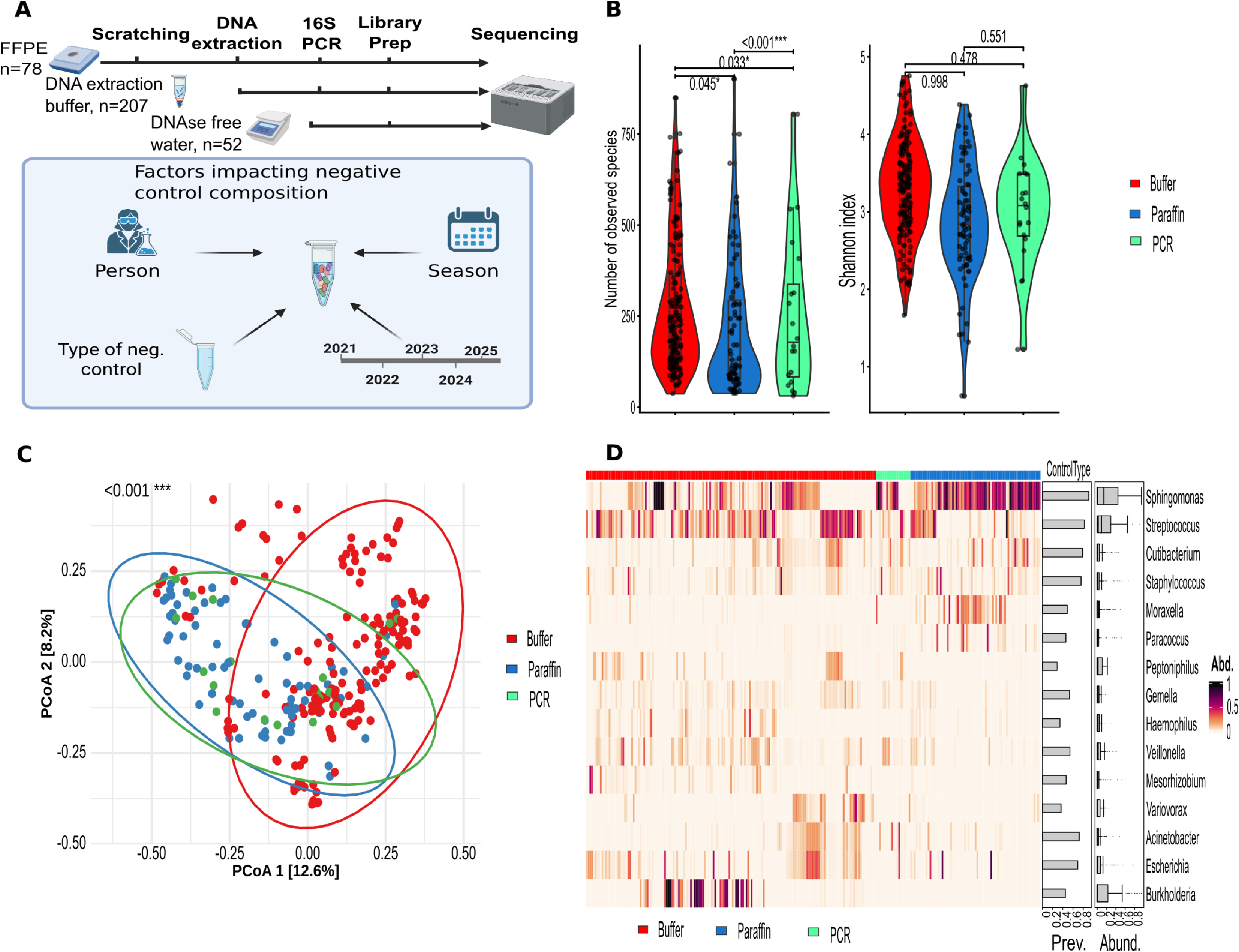
Longitudinal contamination assessment. **A)** Schematic illustration of the collection strategy. Negative controls (NCT) were collected at three steps of the wet-lab workflow together with associated metadata factors including laboratory personnel (LP), sequencing year, and season. **B)** Alpha diversity was analyzed by control type adjusting for LP, year, and season. Observed species were modeled using multivariable negative binomial regression, while the Shannon index was modeled using multivariable gamma regression with a log-link. **C)** Beta diversity analysis of NCT with samples colored and grouped according to control type. P-value derived from distance-based redundancy analyses for Bray-Curtis dissimilarity of control types after adjusting for LP, year, and season. PERMANOVA test was applied **D)** Taxonomic composition heatmap showing the relative abundance of the top 15 genera identified in NCT samples. The overall abundance of each genus across samples is represented by color intensity. Corresponding bar plots summarize the prevalence and abundance distribution of each genus across NCT.

For α diversity, the number of observed species varied significantly among control types (Figure 1B, Table S2). Specifically, the number of distinct observed species in the PCR control was the lowest among three. Moreover, the number of different species in the paraffin control significantly exceeded the ones from buffer controls in multivariable regression analysis controlled for LP, year and season (Table S2). Indeed, a blank paraffin negative control was processed by first scraping paraffin into lysis buffer for DNA extraction, followed by the inclusion of the extracted material in sterile water for subsequent PCR amplification. This sequential workflow (Figure 1A) is consistent with the observed contamination pattern, where paraffin controls harbored the highest number of environmental taxa, followed by buffer and PCR controls. However, no significant differences were observed regarding the Shannon index among different control types, suggesting taxa evenness to play little role among control types. Beta diversity revealed that NCT are significantly clustering with respect to control types (Figure 1C and Table S3). Moreover, NCT were also heavily grouped with respect to LP, year, and season of processing. Indeed, the distance-based redundancy variation was mostly explained by LP (adj. R^2^=0.15), following by year (adj. R^2^=0.07) and season (adj. R^2^=0.06), and last by control type (adj. R^2^=0.02, Table S3).

#### 1.2 Multiple environmental taxa are found in NCTs, at varying abundance

Overall, the NCT contained 856 unique genera, although the community was highly skewed, with the eight most abundant genera accounting for approximately two-thirds of all sequencing reads. These dominant genera included *Sphingomonas*, *Caldibacillus*, and *Streptococcus* (Table S4). In total 3,073 species were detected across all NCT. However, only 110 species reached a relative abundance of ≥0.1%, with a median prevalence of 69 out of 262 samples (Table S5). Next, to explore the biological origin of these bacteria, we focused on dominant genera in our NCT set and revealed a mixture of well characterized environmental microbes and human commensals (Figure 1D). For example, *Sphingomonas*, a ubiquitous environmental genus frequently detected in hospital and laboratory environments, was among the most abundant taxa in nearly all negative controls. In contrast, genera considered as resident microbiota from the oral cavity, like *Haemophilus*, *Streptococcus*, and *Veillonella* (12), were highly prevalent and less abundant in the NCT suggesting an introduction of contamination by LP. This coexistence of environmental and human commensal taxa in NCT requires sophisticated decontamination strategies since an indiscriminate removal of all contamination-associated taxa risks eliminating clinically relevant tumor microbiome signals. Together, our assessment of longitudinally assessed NCT shows a clear impact of a batch effect explained by type of negative control, time of processing and LP.

### 2. Lab personnel have a greater influence on the microbiome than the processing environment

As we discovered highly prevalent and abundant oral microbiota in our NCT and identified LP as major contributor of overall variance in the distance-based redundancy analysis (db-RDA), we further explored the possible batch effect that is introduced by LP that processed the samples. To this end, two LP performed DNA extraction, library preparation and sequencing of four sterile water samples, which were equally distributed. The experiment was repeated in two different environments. First, one LP processed the four negative controls in a low-germ environment and the other one under standard condition. For the second set of negative controls processed by each LP, the LP switched the processing environment (Figure 2A). Notably, α diversity analysis revealed no significant difference with respect to environmental condition (Observed species: IRR = 0.97, p = 0.584, Shannon index: β = 0.95, p = 0.363, Figure S1, Table S6), but a clear distinction between LPs was observed (Observed species: IRR = 1.32, p < 0.001***, Shannon index: β = 1.74, p < 0.001***, Figure 2B and Table S6). In line with these results, β diversity conducted under wrench normalization and Bray-Curtis dissimilarity revealed significant clustering by LP identity (adj R^2^ = 0.46, adj. p-value = 0.005**), with no significant effect of processing environments (Figure 2C, adj R^2^ = 0.05, adj. p-value = 0.245, Table S7). As this study employed a two-factor experimental design, the effect of each factor was evaluated while controlling for the other factor. Across various combinations of distance metrics and abundance transformations, LP remained a significant factor. Furthermore, its consistently high R^2^ values indicate that LP accounts for a substantial proportion of the variation in microbial community composition, highlighting its dominant influence on the sample microbiome profiles. In contrast, the processing environment reached statistical significance only when Bray–Curtis dissimilarity was applied to rarefied data. However, this factor explained only a small proportion of the observed variation (adj. R^2^ = 0.04, adj. p-value = 0.015* vs. LP adj. R^2^ = 0.68, adj. p-value = 0.005**), indicating a comparatively minor effect on microbial community structure (Table S7). Overall, this indicates that while LP-derived contamination is more dominant, environmental setting could contribute an independent, detectable signal.

**Figure 2:**
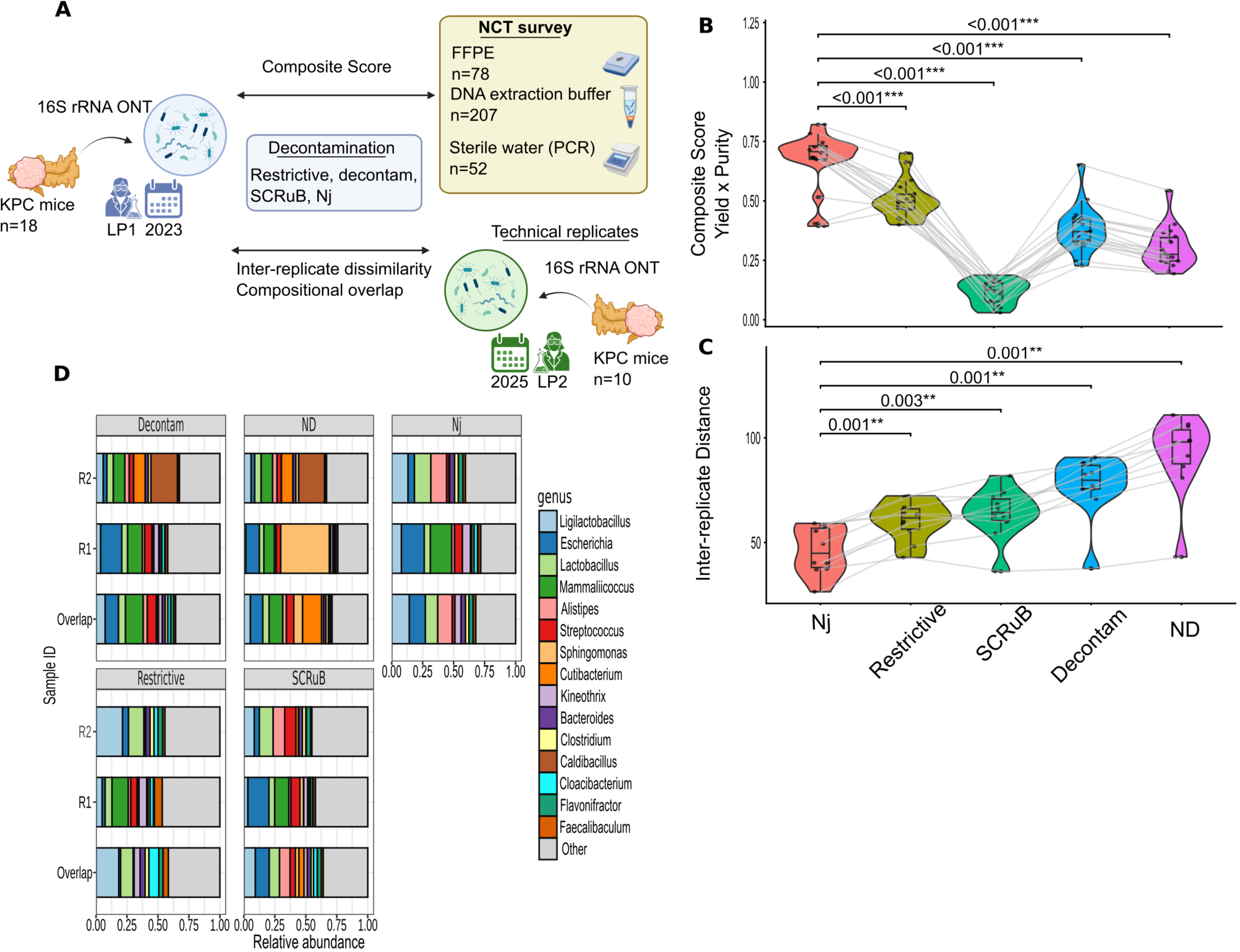
The laboratory personnel (LP) contributes more to the contamination composition than the processing environment. **A)** Illustration of the two-factor experimental design: Each LP processed four shared blank samples within each environment to acquire the lab contaminant profile. Sample processing on the bench wearing standard gloves and no facial mask is referred to as standard environment. Sample processing under the ventilated hood with the LP wearing mask and sterile gloves is referred to as low-germ environment. **B)** Alpha diversity analyses of blank samples, showing the number of observed species and Shannon diversity index of both LP stratified for processing environment. Multivariable negative binomial distribution for number of observed species and multivariable linear regression for Shannon index were applied. **C)** Bray–Curtis dissimilarity for blank samples is displayed colored by LP and shaped by environment. The adjusted p-value was obtained from distance-based redundancy analysis with the LP as explanatory variables controlled for the processing environment. The plot is ordinated with Principal Coordinate Analysis (PCoA).

### 3. A multi-layer statistical testing approach reliably removes contaminants

In light of the low-biomass environment, the assessment of intratumoral microbiota is susceptible to the presence of ubiquitous contaminants. To this end, a thorough decontamination approach is necessary to disentangle contaminants from true signals. To identify the most robust method, we performed benchmarking experiments applying four decontamination strategies, including restrictive filtering, decontam, SCRuB, and multi-layer statistical testing approach (Nj) to full-length 16S rRNA gene profiles of 18 fresh frozen (FF) PDAC samples from *LSL-Kras^G12D/+^*;*LSL-Trp53^R172H/+^*;*Pdx-1-Cre* (KPC) mice bearing endogenous pancreatic tumors that closely recapitulate human PDAC biology and histology (Figure 3A) (13, 14).

**Figure 3:**
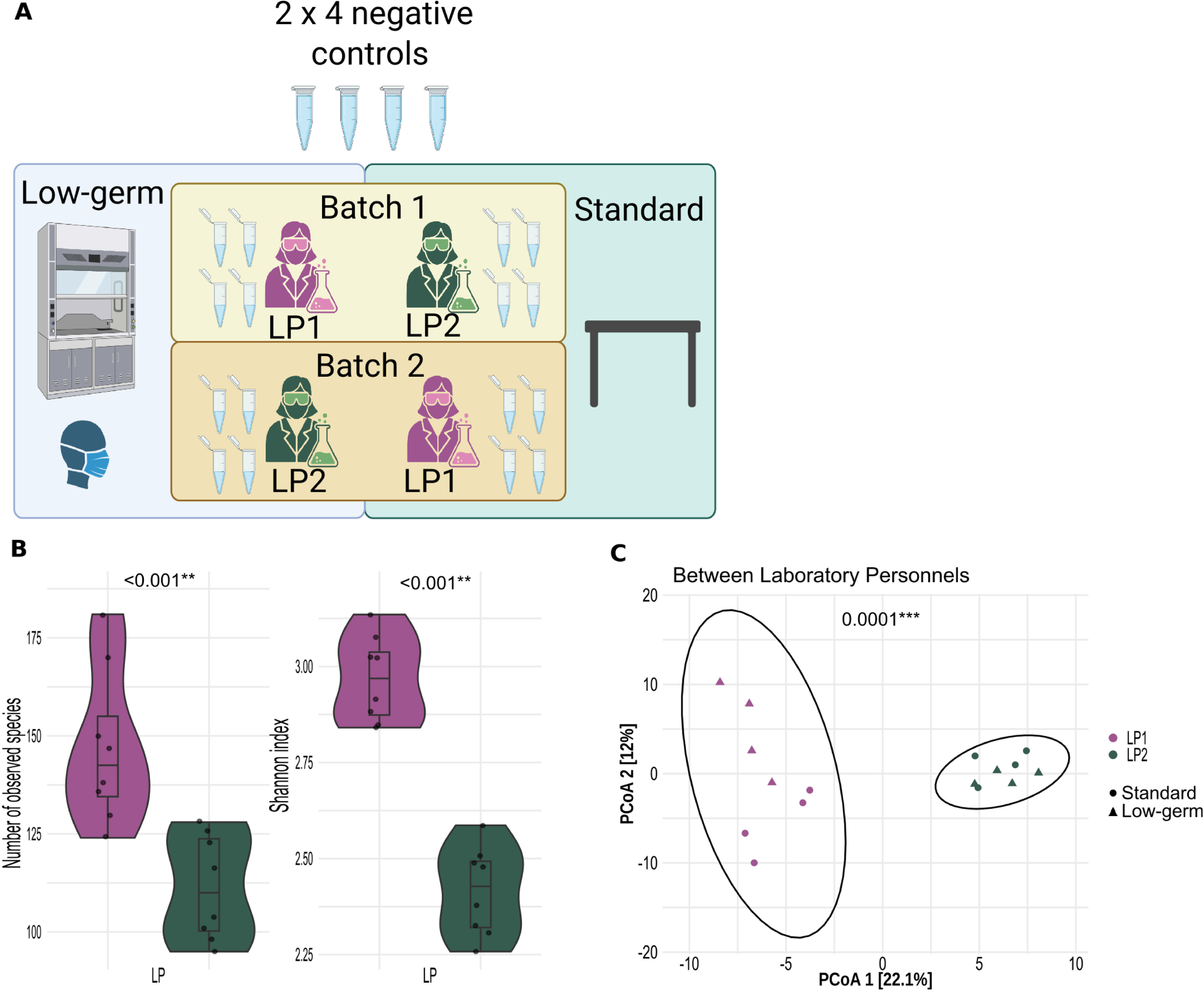
Benchmarking strategy for evaluating microbial decontamination methods. **A)** Experimental design: 18 fresh frozen tumor samples (FF) from *LSL-Kras^G12D/+^*;*LSL-Trp53^R172H/+^*;*Pdx-1-Cre* (KPC) mice were sequenced together with their corresponding negative controls (NCT). Non-decontaminated (ND) profiles were decontaminated through four approaches: restrictive, decontam, SCRuB, and the Nejman et al derived method (Nj) (6). Subsequently, 10 of the 18 FF tumors from KPC mice were re-sequenced as technical replicates using a full-length 16S rRNA gene approach by a different laboratory personnel (LP), employing Oxford Nanopore Technologies (ONT). **B)** Comparison of the four decontamination methods and the ND profile using a paired Wilcoxon signed-rank test based on a composite score integrating sample bacterial profiles and the longitudinal contamination assessment. Higher scores indicate better performance. **C)** Comparison of decontamination methods based on the compositional distance between technical replicates. Inter-replicate dissimilarity was quantified using Aitchison distance of centered-log ratio transformed data, with pairwise comparisons assessed by paired Wilcoxon signed-rank test on species level. Lower inter-replicate distances indicate higher performance. **D)** Overall bacterial composition of the two FF technical replicates and their overlap.

#### 3.1 Assessment Using Longitudinal contamination assessment

In the first approach, we leverage the longitudinal contamination assessment described above as a lab-specific contamination background (CB). Specifically, a bacterial profile, before (non-decontaminated samples, ND) or after decontamination, along with the CB will be used to calculate a corresponding composite score, which rewards the removal of contaminant species in CB while simultaneously preventing the excessive removal of the true taxa.

The Nj method achieved the highest composite score, significantly outperforming all alternatives on species level (Figure 3B, Table S8). ND data scored lower than the Nj, restrictive filtering, and the decontam method. Interestingly, SCRuB achieved the lowest composite score, even significantly smaller than ND.

#### 3.2 Assessment Using Technical Replicates

To provide an orthogonal validation, we re-sequenced 10 FF tumors from KPC mice as technical replicates more than two years after initial sequencing, using a different LP to control for the batch effects. Our underlying assumption was that decontaminated profiles, should exhibit greater concordance between replicates from the same tumor sample than the technical replicates without decontamination (ND).

Inter-replicate dissimilarity was quantified using Aitchison distance of centered-log ratio transformed data, with pairwise comparisons assessed by paired Wilcoxon signed-rank test on species level. The Nj method obtained the smallest inter-replicate distances, indicating superiority within sample consistency (Figure 3C, Table S9). Consistently, inter-replicate among ND samples showed the highest scores, indicating the suspected batch effect. We also validated the results with other distance metrices such as unweighted Jaccard showing a similar pattern (Figure S2).

#### 3.3 Technical replicates with decontamination method reveals reliable intratumoral bacterial profile in fresh-frozen samples from pancreatic ductal adenocarcinoma

Next, we investigated the actual microbial composition after different decontamination approaches to evaluate the biological origin of the respective genera. In the ND profile, samples were dominated by environmental contaminants. *Sphingomonas* predominated in the first replicate of several samples, while *Caldibacillus* (formerly included in *Bacillus*) was prominent in the second replicate of others (Figure 3D). Both genera are considered as environmental taxa rather than mammal commensal bacteria (15, 16). These taxa were largely eliminated following decontamination, though performance varied by methods. The decontam method failed to remove *Caldibacillus* in one replicate, though taking the overlap across replicates substantially reduced its abundance, supporting the use of replicate overlap as a conservative estimate of the true bacterial composition (Figure 3D). A simple taxon-removal strategy based solely on NCT samples, namely the restrictive method, likewise failed to remove contaminants such as *Cloacibacterium* (17). By contrast, Nj and SCRuB yielded similar, more robust compositional profiles. Following Nj decontamination, the majority of remained bacteria belonged to genera with plausible biological relevance to PDAC, including *Ligilactobacillus*, *Escherichia*, *Alistipes*, and *Bacteroides.* All of these are common gut commensals, that may migrate directly into the tumor microenvironment, as described by Schorr et al. 2023 (18).

### 4. Formalin-Fixed Paraffin-Embedded (FFPE) samples are challenging for intratumoral microbiome studies in low-biomass PDAC

To further assess the suitability of FFPE samples for intratumoral microbiome profiling, we evaluated whether microbial signals could be reliably restored from extremely low-biomass PDAC samples. To this end, we examined the concordance of bacterial profiles among paired FFPE and FF samples from ten PDAC samples in the KPC mice cohort (Figure S3A).

For FFPE samples, we compared the original ND profiles and those generated after four decontamination strategies (restrictive filtering, decontam, SCRuB, and Nj) against the bacterial profiles derived from corresponding FF replicates. Notably, regardless of whether the data were ND or processed by any decontamination method, FFPE samples consistently exhibited significantly lower α diversity than the two FF technical replicates, as measured by both the number of observed species and the Shannon index. The two FF technical replicates were highly consistent with each other under most decontamination approaches, with the exception of observed species in the ND and decontam group (Figure 4A).

**Figure 4:**
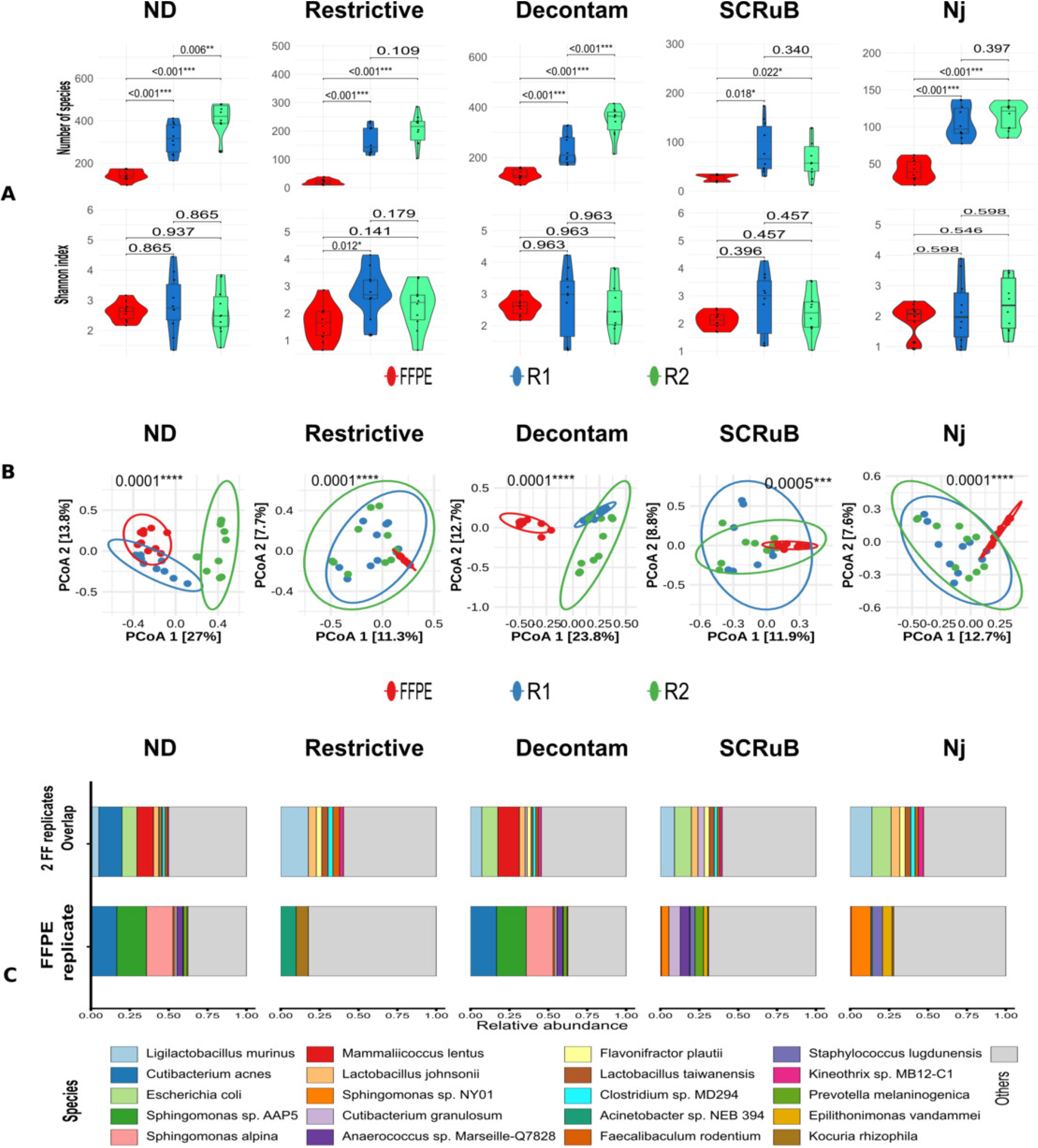
Comparative analysis of paired fresh frozen (FF) and formalin-fixed paraffin embedded (FFPE) tumor samples. **A)** Alpha diversity analyses comparing FFPE samples and the two FF technical replicates across the non-decontaminated (ND) data and four decontamination strategies: restrictive filtering, decontam, SCRuB, and Nejman et al derived method (Nj) (6). Diversity was assessed using the number of observed species and the Shannon index. Univariable negative binomial regression for number of observed species and univariable linear regression for Shannon index were applied. **B)** Bray-Curtis dissimilarity of the two FF technical replicates and paired FFPE samples. Samples are colored and grouped by technical replicates. Analyses were performed using Wrench normalization. The plot is ordinated with Principal Coordinate Analysis (PCoA) and p-values were calculated by PERMANOVA. **C)** Overall bacterial composition of FF overlap and FFPE samples without (ND) and with various decontamination. R – technical replicates

In the db-RDA, Wrench and rarefaction normalization produced comparable results (Figure 4B, S3B). Under both normalization approaches, the ND profile of two FF technical replicates and the corresponding FFPE samples clustered separately. Following decontamination, particularly with the restrictive, SCRuB, and Nj methods, the two FF technical replicates converged into a single cluster, suggesting that decontamination effectively removed sample-specific contaminants and improved inter-replicate concordance (Table S10). In contrast, regardless of the decontamination strategy applied, FFPE samples consistently formed a distinct cluster, indicating that they retained a bacterial profile that differed substantially from their paired FF counterparts (Figure 4B, S3B).

The overall bacterial composition further supported these observations (Figure 4C). Species from genera *Sphingomonas* and *Cutibacterium* remained dominant taxa in both the ND and decontaminated FFPE profiles. In the ND dataset, each sample exhibited a small but detectable overlap in bacterial species shared between the FFPE sample and its two paired FF technical replicates. Although restrictive filtering, SCRuB, and Nj substantially reduced the abundance of these dominant genera, the resulting FFPE microbial communities still showed only minimal compositional concordance with their paired FF samples (Figure S4).

Collectively, these results indicate that even when several NCT and rigorous decontamination strategies are incorporated the authentic intratumoral microbial signal in FFPE samples is difficult to recover from extremely low-biomass PDAC specimens. Residual contaminant taxa still dominate the bacterial profiles, indicating that the decontamination methods applied were unable to reliably preserve the true microbial composition in FFPE samples.

## Discussion

To the best of our knowledge, we are presenting the most extensive longitudinal contamination assessment to date exploring potential factors that contribute to the batch effect. Several studies investigated the ubiquitous bacterial contamination from laboratory reagents (15, 16, 19). Similarly, published dominant genera were also found in our contamination assessment with high prevalence and abundances derived from environmental (e.g. *Sphingomonas*, *Ralstonia, Bradyrhizobium*) and human niches (e.g. *Streptococcus*) (15, 16). Weyrich et al. also conducted a longitudinal assessment including 144 NCT from different sources (16). In line with our findings, they also identified an impact of LP, year and season on the microbial composition. As the laboratory kits are well-known sources of contaminations (15, 19), the human batch effect remains understudied so far. Indeed, the full-length 16S rRNA sequencing of four sterile water samples handled by two LP revealed a strong human effect that significantly outweighed the influence of the processing environment (low-germ vs. standard). Current guidelines for processing and analyzing low-biomass samples do not adequately account for batch effects introduced by different LP (20, 21).

Long-read sequencing has become increasingly interesting for intratumoral microbiome studies because it enables sequencing of full-length 16S rRNA gene amplicons, providing substantially higher taxonomic resolution on species and strain level than short-read approaches (22). This advantage is particularly valuable for low-biomass tumor samples, where limited microbial DNA and high host contamination make accurate taxonomic assignment challenging, particularly for metagenomic sequencing. While short-read 16S rRNA gene sequencing often only resolves microbial communities to the genus level and may fail to distinguish closely related species, full-length 16S rRNA sequencing has demonstrated reliable species and strain level resolution (10, 11, 22). Improved taxonomic precision reduces ambiguity in microbial identification and facilitates more accurate detection of clinically and biologically relevant taxa. This is of particular significance in the context of strain-specific effects of intratumoral microbes (23). Reliable species-level analysis are also possible for 16S rRNA sequencing with short-read technologies by applying multiplexed PCR for several V regions (6). In order to reduce PCR bias and complexity in the wet-lab and bioinformatic pipeline, it appears that a full-length 16S rRNA gene sequencing is a more viable and convenient option (10).

In recent years, a number of highly regarded publications have postulated an impact of intratumoral bacteria on progression, immunosuppression and the determination of subtypes of PDAC (3, 5, 7). However, the revealed intratumoral composition consists mainly of genera like *Pseudomonas*, *Elizabethkingia*, *Sphingobacteria*, *Streptomyces*, *Pseudoxanthomonas*, *Saccharopolyspora*, *Sphingopyxis*. These taxa seem to be rather environmental genera than human commensals. Critically, the methods applied involved no - or only superficial - NCT-controlled decontamination approaches. Despite the compelling mechanistic experiments conducted in these studies, a re-evaluation of the microbial composition detected in tumor samples is necessary. Nejman et al. implemented a thorough and multi-level filtering of contaminants including multiple NCT (6). This approach identified human commensals from the intestinal tract in PDAC samples, thereby providing evidence of the highly probable existence of an intratumoral microbiome. Nevertheless, incorporating hundreds of NCT is largely impractical for mechanism-focused studies and experimental mouse models (13). Furthermore, the applicability of the Nj approach to smaller sample sets has not previously been established. To the best of our knowledge, this study is the first to systematically benchmark different decontamination strategies for PDAC microbiome samples.

Notably, the Nj method consistently outperformed the restrictive filtering, decontam, and SCRuB across all evaluation metrics. This advantage likely stems from Nj’s incorporation of both prevalence-based and rigorous statistical testing, as well as its conservative approach to taxon removal (6). The restrictive decontamination approach, involving the filtration of all species detected in NCT from true samples, is likely to result in the removal of true species from the samples. This is due to the detection of multiple human commensals in the longitudinal contamination assessment, which are highly prevalent (e.g. *Streptococcus oralis* and *Escherichia coli*, Table S5). Decontam is a widely applied tool for contaminant removal. Nevertheless, the authors discussed that the performance is reduced in scenarios, in which the contamination is much more dominant than the true taxa (8). However, the developers of decontam recommend a threshold of 0.5 in these cases, which was applied across all analyses. SCRuB’s relatively modest to poor performance under our assessments may arise from a mismatch between its mechanism and the metric itself. Rather than entirely eliminating contaminant taxa, SCRuB probabilistically subtracts reads attributable to a pooled background source, leaving residual abundances of contaminant taxa. Consequently, a binary presence/absence based metric cannot be applicable, since a down-weighted but non-zero taxon still registers as present. Additionally, performance may be further limited by the absence of well-to-well spatial (plate position) information in our study, as SCRuB is reported to achieve its best results when such information is available. Nevertheless, the overall bacterial composition revealed by SCRuB remained comparable to that of Nj, suggesting that despite its apparent underperformance on our metrics, the method still captured a biologically meaningful signal.

The original Nj decontamination approach was modified for the presented data set. Notably, due to the limited number of NCT (12-15 per batch), binomial statistical tests that were used in the original Nj approach may lack the sensitivity required to effectively filter out false positive contaminant taxa (6). For instance, a species appearing in only a single tumor sample, but absent from the NCT, would pass the binomial test and be misclassified as a true intratumoral taxon. This highlights a potential limitation in contamination detection when NCT sample sizes are insufficient to capture rare contaminants. Thus, instead the Fisher’s exact test was used in our study.

Indeed, PDAC samples decontaminated with the Nj approach harbored several high abundant species in FF that are well-known intestinal commensals from genera such as *Lactobacillus*, *Escherichia*, *Alistipes*, and *Bacteroides*. A similar composition was found in previously analyzed KPC tumor specimen (13). These findings support the presence of a low-biomass microbiome in PDAC and suggest this approach for future intratumoral microbiome studies.

Nevertheless, a significant proportion of species was identified in both replicates following decontamination with the Nj approach, that were not shared between the technical controls (Figure S4). Here, a spatial distribution of microbes has been discussed and observed in several tumor entities (24). Moreover, a PCR bias could result in detection of low-abundant species (20). Finally, the persistence of certain environmental taxa—likely not fully removed by decontamination—highlights the inherent difficulty in assessing a reliable intratumoral microbiome.

The minimal overlap between FF and FFPE samples indicated significant obstacles in the investigation of the intratumoral microbiome using FFPE samples. Nejman et al. were able to detect a reliable microbial composition in FFPE samples (6). However, that was only possible by more than 800 NCT, which is again mostly not feasible for mechanism-focused studies and experimental mouse models.

In summary, our findings from one of the largest longitudinal contaminations assessment underscore the significant human contribution to batch effects, which must be accounted for in low-biomass studies. Moreover, we show that an adjusted Nj approach reliably captures intratumoral microbial composition in FF samples, but not in FFPE tissues, making it suitable for mechanism-based analyses.

## Materials and Methods

### Sample Collection and Processing

NCTs were collected at several steps of a pipeline to account for contamination (Figure 1A). Both NCT and tumor samples were sequenced via full-length 16S rRNA gene. Dorado (version 0.9) was used as a base-caller with q-score cutoff threshold at 9 and MetaPONT (10) was used to obtain bacterial profile of a sample. MetaPONT’s library comprises the complete genome of 13550 bacterial taxa, latest updated in December 2024. Notably, the alignment score and coverage thresholds in MetaPONT were set to 1250 and 70% for tumor samples, and 1000 and 50% for NCT samples, respectively. We used more relaxed cutoff values for NCT samples because they contained a lower amount of bacterial DNA compared with the tumor samples.

Mice experiments were conducted in the Central Animal Experimental Facility at the University Medical Center Goettingen, Germany. Approval was obtained by the LAVES Niedersachsen, Germany (application number 19/3085). Animals were housed under the following conditions: light-dark-cycle of 12 h/12 h, 23 ± 1 °C room temperature (RT), 40–60% humidity, and non-sterile cages. The KPC (*LSL-Kras^G12D/+^*;*LSL-Trp53^R172H/+^*;*Pdx-1-Cre*) mouse model used in this study has been previously described (25). After tumor onset animals were screened 2-3 times a week and sacrificed when the following criteria were fulfilled: overall morbidity, lethargy, signs of pain, rough fur, loss of self-care and social behavior, ascites, cachexia or body weight loss of ≥20% (13). 18 FF from KPC mice were sequenced, together with their corresponding NCT. Later 10 out of 18 FF tumors from KPC mice were sequenced again as technical replicates by another LP.

### Microbiome analysis

DNA isolation, library preparation and sequencing was processed as described previously (10). In brief, PureLink Microbiome kit (Invitrogen) was applied for DNA extraction. The original manufacture’s protocol was adjusted according to the international human microbiome standards (10). Next, a DNA purification was done with a OneStep PCR Inhibitor removal kit (Zymo Research). Full-length16S rRNA gene sequencing was performed with ONT applying the GridION with R10 Flow cells. PCR and library preparation was conducted according to manufacturer’s protocols (ONT). The sequencing process was managed using MinKNOW v.22.08.3.

### Normalization

In the section of the longitudinal contamination assessment as well as contamination investigation in two environments, we utilized taxa_filter function from R package microViz (26) to eliminate extremely rare and possibly noise taxa. The pseudo-code is provided in the technical supplementary file.

In the murine PDAC FF study which comprises PDAC and corresponding negative control samples, we applied slightly different threshold for each category. For tumor samples, we set the threshold for minimal total abundance as 1*10^−6^, while for their respectively negative control we reduce this threshold to 5*10^−7^, due to the assumption that the microbiome content in NCT is much sparer than in tumor sample. Detail and pseudo-code could be found in the technical supplementary file.

We applied rarefaction with even samples library size (for the rarefaction curve, see details in Figure S5-7) and Wrench method (27) in R package for normalization. In the longitudinal contamination assessment, we applied Wrench normalization with respect to control type, whereas in the LP-environment investigation, LP was used as the grouping factor. For the benchmarking of FFPE and FF replicates, Wrench normalization was applied with respect to replicate.

### Decontamination Methods

Four decontamination approaches were evaluated: (i) Restrictive filtering: taxa detected in any NCT samples were excluded from further analyses; (ii) Decontam (8): R package version 1.32.0, with a prevalence-based method classification of contaminant taxa using negative control and p-value cutoff 5; (iii) SCRuB (9): Statistical Correction of Reagent-derived Bias, models contamination as a mixture of biological signal and background noise contaminants and estimates the true microbial composition of each sample.; and (iv) Nejman et. al. procedure (6): the decontamination strategy incorporating multiple filtering criteria. Our adaptation of Nj procedure is as follows. First, taxa whose prevalence is high in NCT samples will be removed. The cutoff threshold was determined based on the relationship between the prevalence cutoff in NCT samples and the proportion of taxa removed from true samples. The selected threshold was required to minimize the removal of taxa from true samples while effectively filtering prevalent contaminants (Figure S8). Second, for those taxa whose prevalence are small in both NCT and true samples, we applied statistical Fisher exact test for each batch to determine if a taxon is a contaminant. The implementation of those four decontamination methods is available in the owncloud repository (https://owncloud.gwdg.de/index.php/s/RPC3bGXwuxlUMHW).

### Composite Score

The composite score was calculated from the bacterial profile before or after decontamination and the lab-specific contamination background (CB) assessed before longitudinally. This score comprised two parts: Yield and purity. Yield is defined as the number of putative true taxa which do not overlap with the CB over the numbers of the total observed taxa. Purity is defined as the fraction between the number of putative true taxa which do not overlap with the CB over the number of putative true taxa from a certain decontamination method. The technical details of the calculation can be found in the Supplementary Methods.

### Statistical Analysis

In the longitudinal contamination assessment and NCT sequencing in different environments by two LP, we applied multivariable generalized models for α diversity metrics and db-RDA (capscale function) for β analysis respectively. Observed species represents overdispersed data, thus a negative binomial regression was applied. For Shannon index a linear or gamma model with log-link was utilized according to the normality. Normal distribution was assessed by Shapiro-Wilk test. Multiple testing correction was performed using Benjamini-Hochberg.

## Data and Code Availability

Fastq files and corresponding metadata and preparation data are available in qiita (Study ID 16568). Relevant R codes are can be freely downloaded from the owncloud repository (https://owncloud.gwdg.de/index.php/s/RPC3bGXwuxlUMHW).

## Abbreviations

DAA: Differential abundance analysis
FF: Fresh frozen samples
CB: Contamination background
Db-RDA: distance-based redundancy analysis
FFPE: Formalin-fixed paraffin-embedded samples
LP: Laboratory person
NCT: Negative controls
ND: Non-decontaminated
Nj: Nejman approach
ONT: Oxford Nanopore Technologies
PDAC: Pancreatic ductal adenocarcinoma
SCRuB: Source-tracking for Contamination Removal in microBiomes

## Acknowledgment

We would like to thank Klara König and Tabea Wollborn for bringing structure and order to the large library of negative controls. All sequencing costs were covered by intramural funding of University Medical Center Goettingen, Forschungsförderungsprogramm 2021 – Startförderung Klinische Studien. The funders had no role in study design, data collection and interpretation, or the decision to submit the work for publication.

## References

1. Yao Y, Zhu Y, Chen K, Chen J, Li Y, Li D, Wei P. 2026. Microbiota in cancer: current understandings and future perspectives. Signal Transduct Target Ther 11:39. doi:10.1038/s41392-025-02335-3.

2. Hanahan D. 2022. Hallmarks of Cancer: New Dimensions. Cancer Discov 12:31–46. doi:10.1158/2159-8290.CD-21-1059.

3. Pushalkar S, Hundeyin M, Daley D, Zambirinis CP, Kurz E, Mishra A, Mohan N, Aykut B, Usyk M, Torres LE, Werba G, Zhang K, Guo Y, Li Q, Akkad N, Lall S, Wadowski B, Gutierrez J, Kochen Rossi JA, Herzog JW, Diskin B, Torres-Hernandez A, Leinwand J, Wang W, Taunk PS, Savadkar S, Janal M, Saxena A, Li X, Cohen D, Sartor RB, Saxena D, Miller G. 2018. The Pancreatic Cancer Microbiome Promotes Oncogenesis by Induction of Innate and Adaptive Immune Suppression. Cancer Discov 8:403–416. doi:10.1158/2159-8290.CD-17-1134.

4. Geller LT, Barzily-Rokni M, Danino T, Jonas OH, Shental N, Nejman D, Gavert N, Zwang Y, Cooper ZA, Shee K, Thaiss CA, Reuben A, Livny J, Avraham R, Frederick DT, Ligorio M, Chatman K, Johnston SE, Mosher CM, Brandis A, Fuks G, Gurbatri C, Gopalakrishnan V, Kim M, Hurd MW, Katz M, Fleming J, Maitra A, Smith DA, Skalak M, Bu J, Michaud M, Trauger SA, Barshack I, Golan T, Sandbank J, Flaherty KT, Mandinova A, Garrett WS, Thayer SP, Ferrone CR, Huttenhower C, Bhatia SN, Gevers D, Wargo JA, Golub TR, Straussman R. 2017. Potential role of intratumor bacteria in mediating tumor resistance to the chemotherapeutic drug gemcitabine. Science 357:1156–1160. doi:10.1126/science.aah5043.

5. Riquelme E, Zhang Y, Zhang L, Montiel M, Zoltan M, Dong W, Quesada P, Sahin I, Chandra V, San Lucas A, Scheet P, Xu H, Hanash SM, Feng L, Burks JK, Do K-A, Peterson CB, Nejman D, Tzeng C-WD, Kim MP, Sears CL, Ajami N, Petrosino J, Wood LD, Maitra A, Straussman R, Katz M, White JR, Jenq R, Wargo J, McAllister F. 2019. Tumor Microbiome Diversity and Composition Influence Pancreatic Cancer Outcomes. Cell 178:795–806.e12. doi:10.1016/j.cell.2019.07.008.

6. Nejman D, Livyatan I, Fuks G, Gavert N, Zwang Y, Geller LT, Rotter-Maskowitz A, Weiser R, Mallel G, Gigi E, Meltser A, Douglas GM, Kamer I, Gopalakrishnan V, Dadosh T, Levin-Zaidman S, Avnet S, Atlan T, Cooper ZA, Arora R, Cogdill AP, Khan MAW, Ologun G, Bussi Y, Weinberger A, Lotan-Pompan M, Golani O, Perry G, Rokah M, Bahar-Shany K, Rozeman EA, Blank CU, Ronai A, Shaoul R, Amit A, Dorfman T, Kremer R, Cohen ZR, Harnof S, Siegal T, Yehuda-Shnaidman E, Gal-Yam EN, Shapira H, Baldini N, Langille MGI, Ben-Nun A, Kaufman B, Nissan A, Golan T, Dadiani M, Levanon K, Bar J, Yust-Katz S, Barshack I, Peeper DS, Raz DJ, Segal E, Wargo JA, Sandbank J, Shental N, Straussman R. 2020. The human tumor microbiome is composed of tumor type-specific intracellular bacteria. Science 368:973–980. doi:10.1126/science.aay9189.

7. Guo W, Zhang Y, Guo S, Mei Z, Liao H, Dong H, Wu K, Ye H, Zhang Y, Zhu Y, Lang J, Hu L, Jin G, Kong X. 2021. Tumor microbiome contributes to an aggressive phenotype in the basal-like subtype of pancreatic cancer. Commun Biol 4:1019. doi:10.1038/s42003-021-02557-5.

8. Davis NM, Proctor DM, Holmes SP, Relman DA, Callahan BJ. 2018. Simple statistical identification and removal of contaminant sequences in marker-gene and metagenomics data. Microbiome 6:226. doi:10.1186/s40168-018-0605-2.

9. Austin GI, Park H, Meydan Y, Seeram D, Sezin T, Lou YC, Firek BA, Morowitz MJ, Banfield JF, Christiano AM, Pe’er I, Uhlemann A-C, Shenhav L, Korem T. 2023. Contamination source modeling with SCRuB improves cancer phenotype prediction from microbiome data. Nat Biotechnol 41:1820–1828. doi:10.1038/s41587-023-01696-w.

10. Ammer-Herrmenau C, Pfisterer N, van den Berg T, Gavrilova I, Amanzada A, Singh SK, Khalil A, Alili R, Belda E, Clement K, Abd El Wahed A, Gady EE, Haubrock M, Beißbarth T, Ellenrieder V, Neesse A. 2021. Comprehensive Wet-Bench and Bioinformatics Workflow for Complex Microbiota Using Oxford Nanopore Technologies. mSystems 6:e0075021. doi:10.1128/mSystems.00750-21.

11. Zhang T, Li H, Ma S, Cao J, Liao H, Huang Q, Chen W. 2023. The newest Oxford Nanopore R10.4.1 full-length 16S rRNA sequencing enables the accurate resolution of species-level microbial community profiling. Appl Environ Microbiol 89:e0060523. doi:10.1128/aem.00605-23.

12. Peng X, Cheng L, You Y, Tang C, Ren B, Li Y, Xu X, Zhou X. 2022. Oral microbiota in human systematic diseases. Int J Oral Sci 14:14. doi:10.1038/s41368-022-00163-7.

13. Pfisterer N, Ammer-Herrmenau C, Antweiler K, Küffer S, Ellenrieder V, Neesse A. 2023. Dynamics of intestinal and intratumoral microbiome signatures in genetically engineered mice and human pancreatic ductal adenocarcinoma. Pancreatology 23:663–673. doi:10.1016/j.pan.2023.07.008.

14. Lee JW, Komar CA, Bengsch F, Graham K, Beatty GL. 2016. Genetically Engineered Mouse Models of Pancreatic Cancer: The KPC Model (LSL-Kras(G12D/+) ;LSL-Trp53(R172H/+) ;Pdx-1-Cre), Its Variants, and Their Application in Immuno-oncology Drug Discovery. Curr Protoc Pharmacol 73:14.39.1–14.39.20. doi:10.1002/cpph.2.

15. Salter SJ, Cox MJ, Turek EM, Calus ST, Cookson WO, Moffatt MF, Turner P, Parkhill J, Loman NJ, Walker AW. 2014. Reagent and laboratory contamination can critically impact sequence-based microbiome analyses. BMC Biol 12:87. doi:10.1186/s12915-014-0087-z.

16. Weyrich LS, Farrer AG, Eisenhofer R, Arriola LA, Young J, Selway CA, Handsley-Davis M, Adler CJ, Breen J, Cooper A. 2019. Laboratory contamination over time during low-biomass sample analysis. Mol Ecol Resour 19:982–996. doi:10.1111/1755-0998.13011.

17. Allen TD, Lawson PA, Collins MD, Falsen E, Tanner RS. 2006. Cloacibacterium normanense gen. nov., sp. nov., a novel bacterium in the family Flavobacteriaceae isolated from municipal wastewater. Int J Syst Evol Microbiol 56:1311–1316. doi:10.1099/ijs.0.64218-0.

18. Schorr L, Mathies M, Elinav E, Puschhof J. 2023. Intracellular bacteria in cancer-prospects and debates. NPJ Biofilms Microbiomes 9:76. doi:10.1038/s41522-023-00446-9.

19. Stinson LF, Keelan JA, Payne MS. 2019. Identification and removal of contaminating microbial DNA from PCR reagents: impact on low-biomass microbiome analyses. Lett Appl Microbiol 68:2–8. doi:10.1111/lam.13091.

20. Fierer N, Leung PM, Lappan R, Eisenhofer R, Ricci F, Holland SI, Dragone N, Blackall LL, Dong X, Dorador C, Ferrari BC, Goordial J, Holmes SP, Inagaki F, Korem T, Li SS, Makhalanyane TP, Metcalf JL, Nagarajan N, Orsi WD, Shanahan ER, Walker AW, Weyrich LS, Gilbert JA, Willis AD, Callahan BJ, Shade A, Parkhill J, Banfield JF, Greening C. 2025. Guidelines for preventing and reporting contamination in low-biomass microbiome studies. Nat Microbiol 10:1570–1580. doi:10.1038/s41564-025-02035-2.

21. Eisenhofer R, Minich JJ, Marotz C, Cooper A, Knight R, Weyrich LS. 2019. Contamination in Low Microbial Biomass Microbiome Studies: Issues and Recommendations. Trends Microbiol 27:105–117. doi:10.1016/j.tim.2018.11.003.

22. Johnson JS, Spakowicz DJ, Hong B-Y, Petersen LM, Demkowicz P, Chen L, Leopold SR, Hanson BM, Agresta HO, Gerstein M, Sodergren E, Weinstock GM. 2019. Evaluation of 16S rRNA gene sequencing for species and strain-level microbiome analysis. Nat Commun 10:5029. doi:10.1038/s41467-019-13036-1.

23. Zepeda-Rivera M, Minot SS, Bouzek H, Wu H, Blanco-Míguez A, Manghi P, Jones DS, LaCourse KD, Wu Y, McMahon EF, Park S-N, Lim YK, Kempchinsky AG, Willis AD, Cotton SL, Yost SC, Sicinska E, Kook J-K, Dewhirst FE, Segata N, Bullman S, Johnston CD. 2024. A distinct Fusobacterium nucleatum clade dominates the colorectal cancer niche. Nature 628:424–432. doi:10.1038/s41586-024-07182-w.

24. Galeano Niño JL, Wu H, LaCourse KD, Kempchinsky AG, Baryiames A, Barber B, Futran N, Houlton J, Sather C, Sicinska E, Taylor A, Minot SS, Johnston CD, Bullman S. 2022. Effect of the intratumoral microbiota on spatial and cellular heterogeneity in cancer. Nature 611:810–817. doi:10.1038/s41586-022-05435-0.

25. Hingorani SR, Wang L, Multani AS, Combs C, Deramaudt TB, Hruban RH, Rustgi AK, Chang S, Tuveson DA. 2005. Trp53R172H and KrasG12D cooperate to promote chromosomal instability and widely metastatic pancreatic ductal adenocarcinoma in mice. Cancer Cell 7:469–483. doi:10.1016/j.ccr.2005.04.023.

26. Barnett D, Arts I, Penders J. 2021. microViz: an R package for microbiome data visualization and statistics. JOSS 6:3201. doi:10.21105/joss.03201.

27. Kumar MS, Slud EV, Okrah K, Hicks SC, Hannenhalli S, Corrada Bravo H. 2018. Analysis and correction of compositional bias in sparse sequencing count data. BMC Genomics 19:799. doi:10.1186/s12864-018-5160-5.

